# Altered Stress and Fear Responses in the VPA Rat Model of Autism: Behavioral Dissociation Across Tactile, Nociceptive, and Social Contexts

**DOI:** 10.1101/2025.07.07.663538

**Authors:** Debora Hashiguchi, Ana Luiza Dias, Rodrigo Neves Romcy-Pereira

## Abstract

Sensitivity to environmental stimuli is a fundamental aspect of human behavior, and its dysregulation is associated with stress-related and anxiety disorders. In autism spectrum disorder (ASD), altered sensory processing may contribute to increased vulnerability to such conditions. To better understand this relationship, we evaluated autonomic and behavioral responses to tactile, nociceptive, and social stressors in juvenile Wistar rats prenatally exposed to valproic acid (VPA), a widely used animal model of ASD. VPA-treated and saline-treated control (CTL) rats underwent a battery of stress-related behavioral tests. Defecation, freezing and vocalization behaviors were analyzed in response to handling, electro-tactile stimulation, nociceptive foot shock (fear conditioning) and social stress stimuli (emotional contagion). VPA-treated rats maintained increased defecation during handling and a higher prevalence of defecation during electro-tactile stimulation, without corresponding changes in freezing. During fear conditioning, these animals showed delayed onset and heightened freezing responses. Furthermore, unlike CTL rats, VPA-treated rats lacked correlations among freezing, defecation, and vocalization. In the emotional contagion paradigm, observation of shock in the partner increased the prevalence of animals freezing and reduced the prevalence of animals vocalizing in both VPA and CTL groups (oV+ and oC+) compared to their respective controls (oV- and oC-). However, these effects were more sustained in oV+ rats, which also exhibited prolonged freezing behavior and an earlier reduction in vocalization rate. These findings indicate that VPA-treated rats display heightened stress reactivity, habituation deficit and disrupted coordination of fear responses, supporting the VPA model as a relevant tool for investigating the neurobiological basis of stress vulnerability and social dysfunction in ASD.

## Introduction

The ability to respond to environmental changes is a fundamental characteristic of human existence. Excessive worry, fear, and heightened sensitivity to threats characterize stress and anxiety disorders. Evidence indicates that individuals with ASD are especially susceptible to experiencing anxiety (1–7). Studies suggest that innocuous stimuli are more frequently appraised as stressful by affected individuals compared to normotypical individuals, which may result in behaviors similar to the characteristics inherent to ASD, such as difficulties in coping with changes, abnormal responsiveness to sensory stimuli and social demands, thus increasing the vulnerability of these individuals to stress disorders (1,8–12).

In humans and animals, the primary arousal system in the human body is the autonomic nervous system (ANS). The ANS responds rapidly to stress by activating and inhibiting its sympathetic and parasympathetic branches. These responses manifest in a variety of physiological changes, including increased heart rate, blood pressure, respiratory rate, pupil dilation, and colonic motility (13–17). These responses collectively prepare the body to confront or avoid the stressor, commonly known as the fight-or-flight response. Behavioral responses to these physiological changes include increased alertness and focus, enhanced sensory perception, and suppression of nonessential functions. Consequently, studies use changes in cardiac activity, sweating, skin temperature, and colonic motility as indicators to assess ANS functionality (18–22). Increased fecal pellet release, driven by increased colonic motility, and freezing behavior, which involves the integration of central nervous system, ANS, and sensorimotor functions, have been observed in stressed rats (23–26). However, the behavioral response elicited varies depending on the level of disturbance in ANS homeostasis. Furthermore, stress-induced emission of ultrasonic vocalizations (USVs) within the frequency range of 20–35 kHz, also known as 22 kHz calls, has been widely correlated with the level of stress experienced by the animal (27,28). Calls lasting less than 100 ms are typically associated with acute distress, while longer calls, which can extend up to 3000 ms, are frequently observed under sustained stressful conditions (29–33). All of these behaviors are recognized as stress signals and considered vital for social communication, particularly in signaling danger within a group.

The valproic acid (VPA) rat model is extensively employed as a preclinical model of autism spectrum disorder (ASD) due to its recapitulation of key neural and behavioral alterations characteristic of the condition (34–37). Despite its widespread application, the model’s validity for investigating stress susceptibility remains underexplored. This study aimed to assess the feasibility of employing the VPA rat model for translational research on stress responses to tactile, nociceptive, and adverse social stimuli. To achieve this, four experimental paradigms were employed: (1) a manipulation test and (2) an electro-tactile sensitivity test to evaluate stress responses to tactile stimuli; (3) a fear conditioning paradigm to assess stress responses to nociceptive stimuli; and (4) an emotional contagion paradigm to investigate stress responses to negative social stimuli.

## Material and methods

### Animals

Experimental and control animals were obtained by breeding male and female Wistar rats from the Brain Institute Animal Facility (BISIC) at the Federal University of Rio Grande do Norte (UFRN). Breeding pairs were >60 days old and weighed between 250–300 g at the time of mating. On gestation day 12.5, 14 pregnant females were injected with either sodium valproate (n = 8; 250 mg/mL saline solution, 500mg/Kg, i.p., VPA; Sigma-Aldrich P-4543) or saline (n = 6; 0.15M NaCl) solutions. A group of pregnant females (n = 9) was left undisturbed for the generation of naive animals to be used in the emotional contagion paradigm (as demonstrators). In total, we used 23 litters generated by mating 17 females with 14 males. Six females were mated twice and contributed with two litters (first litter control and second litter VPA, n = 1; first litter control and second litter naive, n = 1; first litter naive and second litter VPA, n = 3; or first litter VPA and second litter naive, n = 1). After birth, pups were left in their homecage with the dam until weaning at P21.

Newborn rats used in the tactile sensitivity tests were transferred at P21 to new clean cages (3-6 littermates/cage) without sexual segregation. A total of 58 animals were included in these experiments: 28 VPA-treated rats (15 females and 13 males) and 30 CTL rats (15 females and 15 males). Behavioral tests were conducted between P33 and P37. Newborn animals used in the nociceptive and negative social experiments were transferred at P21 to new clean cages in pairs. In each cage, a VPA-treated or CTL rat was housed with a naive rat of the same age and sex. A total of 122 animals were included in these experiments: 31 VPA-treated rats (15 females and 16 males), 30 CTL rats (15 females and 15 males) and 61 naive rats (30 females and 31 males). The same cohorts of VPA and CTL animals were used for the nociceptive and negative social experiments. Behavioral tests were performed on P38 and P42.

All VPA-treated rats showed a characteristic tail kink due to the VPA embryological effect. All experimental procedures used in this study were approved by the Ethics Committee on the Use of Animals (CEUA-UFRN, protocol n° 017/2020). Throughout the experiments, all animals were kept in polypropylene boxes (55 x 46 x 30 cm) in a conditioned environment, with temperature set to 24°C, 70% humidity, a 12h/12h light-dark cycle (lights-on: 7am; lights-off: 7pm), with food and water freely available.

### Behavioral tests

#### Touch sensitivity test

To investigate touch sensitivity, we used gentle handling as a stimulus. Over a period of four days, rats aged P33 to P36 were transferred from the vivarium to an experimental room within their housing boxes. Once in the experimental room, each rat was individually removed from its cage and handled by the experimenter. Handling involved four sessions/day of one minute each, during which the animals received gentle hand strokes on their fur while held in the experimenter’s hand. Between sessions, animals were transferred back to their home cage where they stayed for five minutes before the next session. Handling sessions were carried out at the end of the lights-on phase, between 4pm and 7pm. Once the handling was completed, the animals were returned to the vivarium.

#### Electro-tactile sensitivity test

To investigate electro-tactile sensitivity of the glabrous skin of the paws, we used footshocks as stimuli. The experimental apparatus consisted of a shock chamber measuring 32 cm in length, 25 cm in width, and 30 cm in height, featuring a metallic grid floor for the delivery of the electrical foot-shock stimuli. The chamber had a translucent roof with small openings on the sides, two metallic walls, and two translucent walls, through which video recordings were obtained. Video monitoring captured the entire area of the apparatus, through two high-definition webcams (Logitech C920). Foot-shock stimuli were applied through the grid by a programmable digital stimulus generator (STG 4002 Multi Channel Systems) controlled by software (MC stimulus 4002). Each stimulus consisted of a rectangular pulse of 1 sec duration and varying amplitude (50-550 μA; 50 μA steps). A digital ultrasonic microphone (Ultramic 250K, Dodotronics) was also positioned through the opening at the roof of the chamber to record vocalizations.

Microphone and video camera recordings were synchronized to the foot-shocks by an Arduino device programmed to activate a light flash and a sound buzz simultaneous to the electrical stimulus.

All animals were tested one day after the last handling session, which was used as habituation to the experimenter and to the experimental room. The experiment took place at the end of the lights-on phase, between 4pm and 7pm and was conducted under dim light. For the experiment, the rats were placed in the middle of the apparatus facing one of the metal walls. After a period of 90 sec of free exploration, single electrical stimuli of 1 sec were applied with increasing intensity in 50 µA steps with random inter-stimulus intervals between 30 and 90 sec. The test was considered completed when the animal displayed jump or flee, for two consecutive stimuli.

#### Nociceptive stress response

To investigate the nociceptive stress response, we used a contextual fear conditioning paradigm with repeated unpredictable footshocks. The apparatus consisted of an operant conditioning chamber (32L x 25W x 30H cm each) with a metal grid floor, a translucent roof with small openings on the sides, surrounded by two metallic and two translucent walls. Electrical foot shock stimuli were applied by a programmable digital stimulus generator (STG 4002 MultiChannel Systems) controlled by software (MC stimulus 4002). Video monitoring captured the entire area of the apparatus, through a high- definition webcam (Logitech C920) positioned above the apparatus. A digital ultrasonic microphone (Ultramicrophone 250K, Dodotronic; 16 bits ADC; 250kHz sampling rate) was positioned through the opening at the roof of the chamber to record vocalizations at approximately 30 cm from the chamber floor. Microphone and video camera recordings were synchronized to the foot-shocks by an Arduino device programmed to activate a light flash and a sound buzz simultaneous to the electrical stimulus. Videos recordings in .mp4 format and audio recordings in .wav format were stored on a hard disk for posterior analysis.

The experiment took place at the end of the lights-on phase, between 2pm and 5pm and was carried out under dim light. At P38, rats were placed in the middle of the apparatus facing one of the metal walls. Following a 4-min period of free exploration that served as baseline (block B0), five electrical shock stimuli of 1-sec duration and random inter-stimulus interval 0-180 sec were applied (blocks B1-B5). The experiment finished 4 minutes after the last shock was applied. We used a shock intensity (750uA) that corresponded to two times the nociceptive threshold defined during the electro-tactile sensitivity test. At the end of the experiment, each rat was transferred to a new clean (neutral) cage for one hour before returning to its home cage.

#### Social stress response

In order to investigate the animal’s response to a cagemate stress demonstration, we used the emotional contagion paradigm, in which an experimental animal (observer, OBS) observing a conspecific (demonstrator, DEM) undergo a series of aversive foot shocks. The apparatus used consisted of two compartments: a conditioning chamber (shock chamber) adjacent to a neutral chamber (32L x 25W x 30H cm) was separated by a translucent plexiglass wall with two rows (4 and 6 cm above the floor) of 8 open roles each (1cm diameter). An ultrasonic microphone was positioned in the shock chamber through a roof aperture in the chamber (30 cm from the floor) to record vocalizations. Control software and equipment synchronization were the same as previously described. Audio (.wav files) and video recordings (.mp4 files) were save and stored in hard disk for posterior analysis of freezing and vocal behaviors. Habituation and testing were conducted during the lights-off phase, between 7:00 p.m. and 3:00 a.m., under dim light.

Over the following two days of fear conditioning, OBS animals were habituated to the neutral chamber for 10 minutes each day (P39-P40). They were placed in the center of the neutral box, facing the shock chamber, and allowed to freely explore the apparatus and then, returned to their home cages. At P41, both OBS and DEM were brought to the experimental apparatus. The DEM animal was placed in the center of the shock chamber, facing one of the side walls, while the OBS was placed in the center of the neutral chamber, facing one of the side walls. Following a 4-min period of free exploration (B0), which served as a baseline, five electrical shock stimuli of 1-sec duration, 750 µA intensity, delivered at random inter-stimulus interval 0-360 sec, were applied to the DEM animal (blocks B1-B5). In order to maximize the OBS-to-DEM attention, all shock stimuli were delivered to the DEM only when the OBS animal faced the DEM. Following the last stimulus, both animals remained for an additional 4 min in the apparatus. Throughout the experiment, the OBS animal was able to make visual contact with the DEM through a clear plexiglass division. All experiments were conducted under dim light conditions.

In order to assign the identity of the call emitter while using one microphone in the DEM chamber, we designed a pilot experiment (Fig S1). Vocalizations above -76 dB were considered as originating from animals in the same chamber where the microphone was placed (DEM chamber). Calls with power below -85 dB were considered predominantly originating from animals in the adjacent chamber (OBS chamber). In summary, high intensity vocalizations were considered DEM calls, whereas low intensity vocalizations were considered OBS calls.

Figure S1: **USV power.** (A) Maternal separation (MS) experimental design at P08 and graphic with USVs mean power by animal. Dotted line represents the lower power of USV emitted in the microphone chamber (– 85.84 kHz). Dashed line represents the higher power of USV emitted in the opposite chamber (– 84.03 kHz). (B) Maternal separation (MS) experimental design at P14 and graphic with USVs mean power by animal. Dotted line represents the lower power of USV emitted in the microphone chamber (– 90.08 kHz). Dashed line represents the higher power of USV emitted in the opposite chamber (– 83.60 kHz). (C) Fear conditioning (FC) experimental design at P35-38 and graphic with USVs mean power by animal. Dotted line represents the lower power of USV emitted in the microphone chamber (– 92.12 kHz). Dashed line represents the higher power of USV emitted in the opposite chamber (– 76.54 kHz). ^#^ between-groups significant difference.

### Behavioral analysis

Quantification of fecal pellets was recorded at the end of each handling session and immediately after all other experiments. To maintain objective quantification of variables, video and audio files (.mp4 and .wav) were randomly coded and analyzed by a skilled experimenter.

For the determination of the tactile threshold, the following behaviors were considered: grid investigation and paw retraction. The tactile threshold was defined as the lowest current at which the animal exhibited any of the previous behaviors. For the determination of the nociceptive threshold, the following behaviors were considered: jumping and fleeing. The nociceptive threshold was defined as the lowest current at which the animal displayed any of these behaviors. A freezing episode was identified as the behavioral state where the animal remained completely still without any intentional movement for a duration exceeding 3 seconds. It was quantified as the relative freezing time throughout the experiment. The analyzes of nociceptive and social negative experiments compared measurements during the baseline (B0) with either their combination during all shock blocks (B1-5) or individually with their values per block (B1 to B5). Following B0 (baseline, 4 min before first shock), each subsequent block (B1-B4) extended from 2s prior to a shock to 2s before the next shock. The last block (B5) extended from 2s before the fifth shock to 4 min after this shock. All video recordings were blindly analyzed by behavioral logging using BORIS software v.7.13.8.

The vocal behavior during the experiments was analyzed by quantifying the number of USVs produced and their acoustic properties. The analysis consisted of four steps: (1) Call detection, (2) Curation, (3) Segmentation and (4) Feature extraction. They were performed using software routines developed in our laboratory (Bessa, Rafael dos Santos de & Romcy-Pereira, 2023) and a custom-modified version of DeepSqueak software v2.6.2. Automatic USV detection was performed by computing a time-dependent entropy score of the power distribution across frequencies in the audio spectrogram with time resolution = 2 ms; frequency resolution = 65 Hz; frequency range, 20-100 kHz using the Chronux software package (Bokil et al., 2010). USV curation and segmentation consisted in a software-assisted refinement of USV time and frequency boundary boxes, which were performed by a trained investigator. The software performed a pre-identification of the USVs, which could be confirmed or rejected by the investigator. The investigator could include USVs that were not identified by the software or exclude regions of noise incorrectly identified as USVs. The criteria used to consider a putative USV were as follows: minimum frequency of 20 kHz, maximum frequency of 125 kHz, minimum duration of 5 ms, and a maximum interval between consecutive signals of 30 ms. For the purposes of this study, statistical analysis focused on the number of calls produced, call duration, and fundamental frequency of long 22 kHz calls (< 35-kHz; > 200 ms).

### Statistics

We applied parametric tests when data distributions conformed to normality criteria, otherwise non-parametric tests were used. For proportions and percentages, data were z-transformed before applying a parametric test. Fisher’s exact test or Chi-square were employed to compare data on contingency tables from two independent samples. Mann-Whitney and Student-t tests were employed to compare two independent distributed samples. Paired Student-t test, Wilcoxon signed-rank test employed to compare single group paired samples. Friedman followed by post hoc Dunn’s tests and repeated measures (RM) one-way ANOVA with subsequent Holm-Sidak post-hoc test was employed in single group repeated measures. Kruskal-Wallis was employed to compare multiple groups independent data. Two-way RM ANOVA followed by post-hoc Holm-Sidak test was employed to compare to between-groups with repeated measures. Two-way RM ANOVA with subsequent Dunnets post-hoc test was employed to compare to within-group repeated measures. Within-group analysis was used on data spanning multiple experimental time blocks. Spearman correlation was applied to evaluate linear relationship between behavioral variables. Simple linear regression was applied to evaluate correlations. Correction for multiple tests was applied to original p-values when necessary, by computing a new set of adjusted p-values through the two-stage False Discovery Rate (FDR) proposed by Benjamini, Krieger and Yekutieli (38). Statistical significance level was set at 0.05. Statistical analyses were conducted using GraphPad Prism software (v9.0.0).

## Results

### Increased sensitivity and delayed habituation to repeated handling in VPA-treated rats

Over four handling sessions (Fig 1A), a similar proportion of VPA-treated and CTL animals displayed defecation behavior, defined as the release of at least one fecal pellet (Fig 1B). Analysis across experimental days showed a higher prevalence of VPA-treated animals releasing fecal pellets compared to controls on the second day (Fig 1C). Within-group analysis showed a significant decrease in the relative number of CTL animals defecating after the first session indicating an early habituation to the handling stress. Conversely, such a decrease occurred in the VPA group by the third day onward. We also quantified the number of fecal pellets produced by responsive animals, in order to assess the magnitude of their behavioral responses. Our results showed that there was no overall significant difference in the number of fecal pellets released by CTL and VPA animals during the four experimental days (Fig 1D). However, 33% of VPA-treated rats released more than 10 fecal pellets compared to 6% of CTL rats, indicating higher sensitivity to handling stress among VPA-treated animals. More fecal pellets were released by VPA animals on the second day compared to CTL animals (Fig 1E). Temporal analysis across handling sessions showed that while CTL rats habituated (reduced release of fecal pellets) from second day on, VPA-treated animals showed delayed habituation starting from day 3 on. We also assessed differences between sexes by comparing female and male subjects within each group. No significant differences were observed either in the prevalence or in the magnitude of the defecation response (Data not shown).

**Fig 1.**
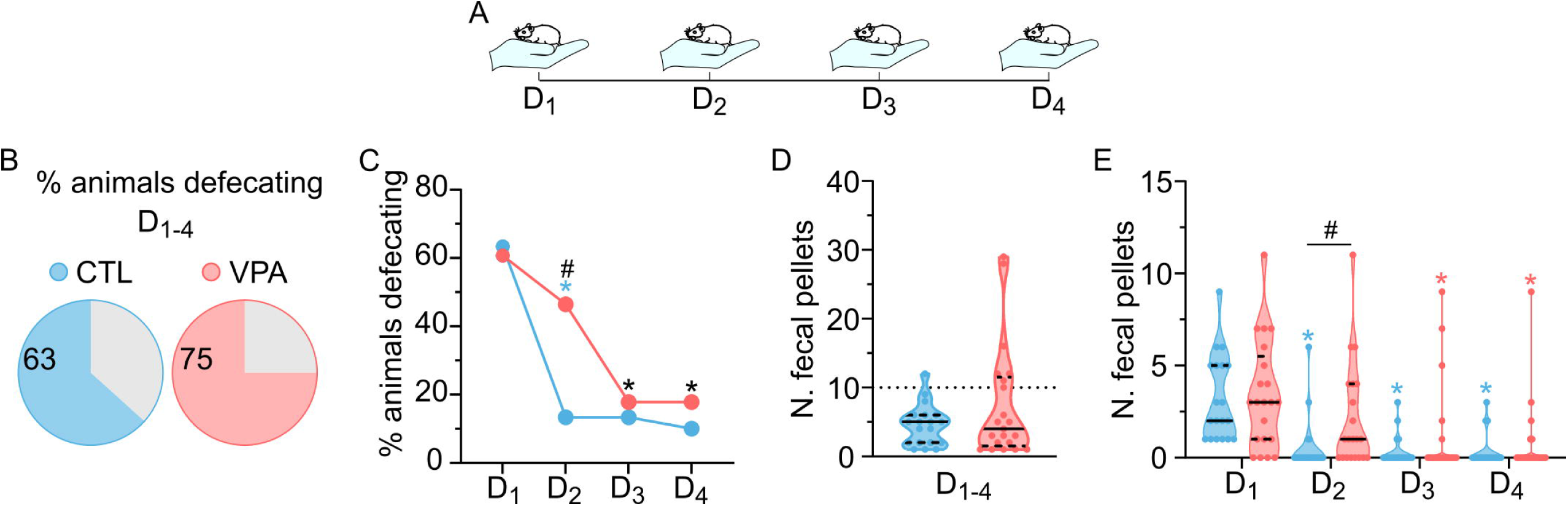
Stress responses elicited by handling. (A) Schematic illustration of the experiment. Four handling sessions of 1 min with 5 min intervals between handlings, once a day for four consecutive days. (B) Percentage of animals releasing fecal pellets over all experimental days. ^#^p > 0.050. (C) Percentage of animals releasing fecal pellets on each experimental day. ^#^p = 0.030; CTL *p = 0.001; VPA *p = 0.008. (D) Number of fecal pellets released over all experiment days. Dotted line for reference at 10 fecal pellets. ^#^p > 0.050. (E) Number of fecal pellets on each experimental day. ^#^p = 0.030; CTL *p ≤ 0.004; VPA *p = 0.008. Panels B and C, Fisher’s exact test [N_CTL_ = 30; 15 female, 15 male. N_VPA_ = 28; 15 female, 13 male]. Panel D, Mann-Whitney test. Panel E, Mann-Whitney and Friedman tests. [N_CTL_ = 19; 10 female, 9 male. N_VPA_ = 21; 10 female, 11 male] ^#^, Between- group significant differences. *, Within-group significant difference compared to D1.

### Increased susceptibility of VPA-treated rats to electro-tactile stimulation

During the electro-tactile sensitivity test (Fig 2A), the VPA group showed a significantly prevalence of animals releasing fecal pellets as compared to the CTL group (Fig 2B). However, the number of pellets released per responsive animal did not differ between VPA and CTL groups (Fig 2C). Analysis of freezing behavior demonstrated no significant group differences in either the prevalence of animals displaying freezing (Fig 2D) or the relative duration of freezing among responsive animals (Fig 2E).

**Fig 2:**
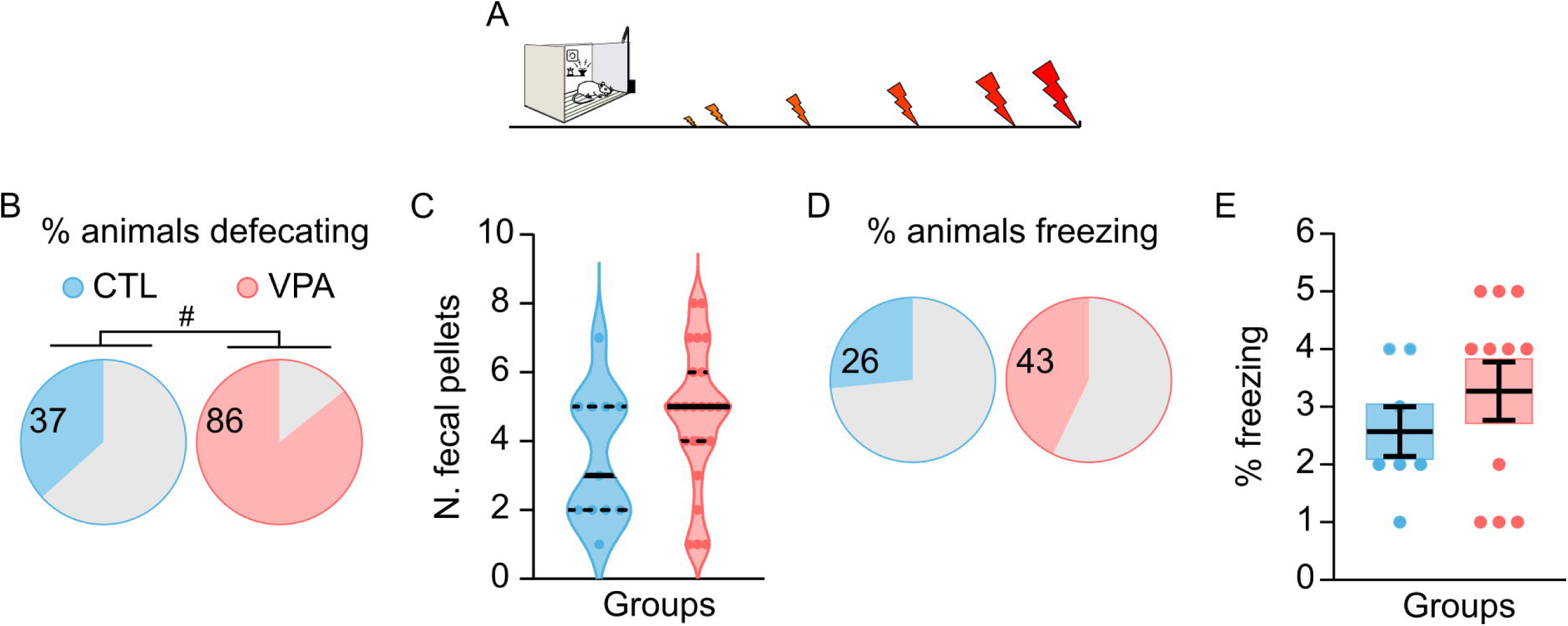
Stress responses elicited by electro-tactile stimuli. (A) Experimental design: the rats were placed in the middle of the apparatus facing one of the metal walls. After a period of 90 sec of free exploration, single electrical stimuli of 1 sec were applied with increasing intensity on 50 µA steps, random inter-stimuli intervals between 30 and 90 sec. The test ended with the presentation of nociceptive variables by two consecutive stimuli. (B) Percentage of animals that released fecal pellets. ^#^p < 0.001. (C) Number of fecal pellets released. ^#^p > 0.050. (D) Percentage of animals displaying freezing behavior. ^#^p > 0.050. (E) Percentage of freezing behavior. ^#^p > 0.050. Panels B and D, Fisher’s exact test [N_CTL_ = 30; 15 female, 15 male. N_VPA_ = 28; 15 female, 13 male]. Panel C, Mann-Whitney [N_CTL_ = 11; 7 female, 4 male. N_VPA_ = 24; 14 female, 10 male]. Panel E, [N_CTL_ = 8; 5 female, 3 male. N_VPA_ = 12, 6 female, 6 male]. ^#^, Between-group significant differences.

Additionally, assessments of tactile and nociceptive thresholds indicated no significant differences between VPA-treated and CTL animals (Fig S2B). In order to assess sex differences, we compared the behavior of females and males in each group. Our results showed no sex differences in stress susceptibility, magnitude of defecation response or freezing behavior in the VPA or CTL group. Similarly, no sex differences were detected in the nociceptive threshold. However, female CTL rats exhibited higher tactile thresholds compared to their male counterparts, whereas no sex differences were observed among VPA-treated animals (Fig S2C).

### Delayed onset and increased magnitude of freezing in VPA-treated rats during fear conditioning

VPA and CTL groups had similar prevalence of animals displaying defecation behavior during fear conditioning (Fig 3A and 3B). No significant difference was detected in the number of fecal pellets produced by responsive rats in either group (Fig 3C). Freezing behavior was also similarly prevalent in CTL and VPA-treated animals (Fig 3D). Within-group analyses revealed that, in both the VPA and CTL groups, the prevalence of freezing behavior significantly increased from the second block (B2) onward compared to baseline (B0; Fig 3E). Comparing the groups, responsive animals differ in their overall freezing behavior (Fig 3F). Initially, VPA-treated rats exhibited a delayed onset of freezing compared to CTL. While CTL animals showed a significant increase in freezing behavior from B1 onward relative to baseline (B0), VPA-treated rats only displayed a comparable increase starting at B2. Moreover, VPA-treated animals exhibited a more pronounced freezing response, peaking at B4, where a statistically significant difference was observed compared to CTL animals.

**Figure 3:**
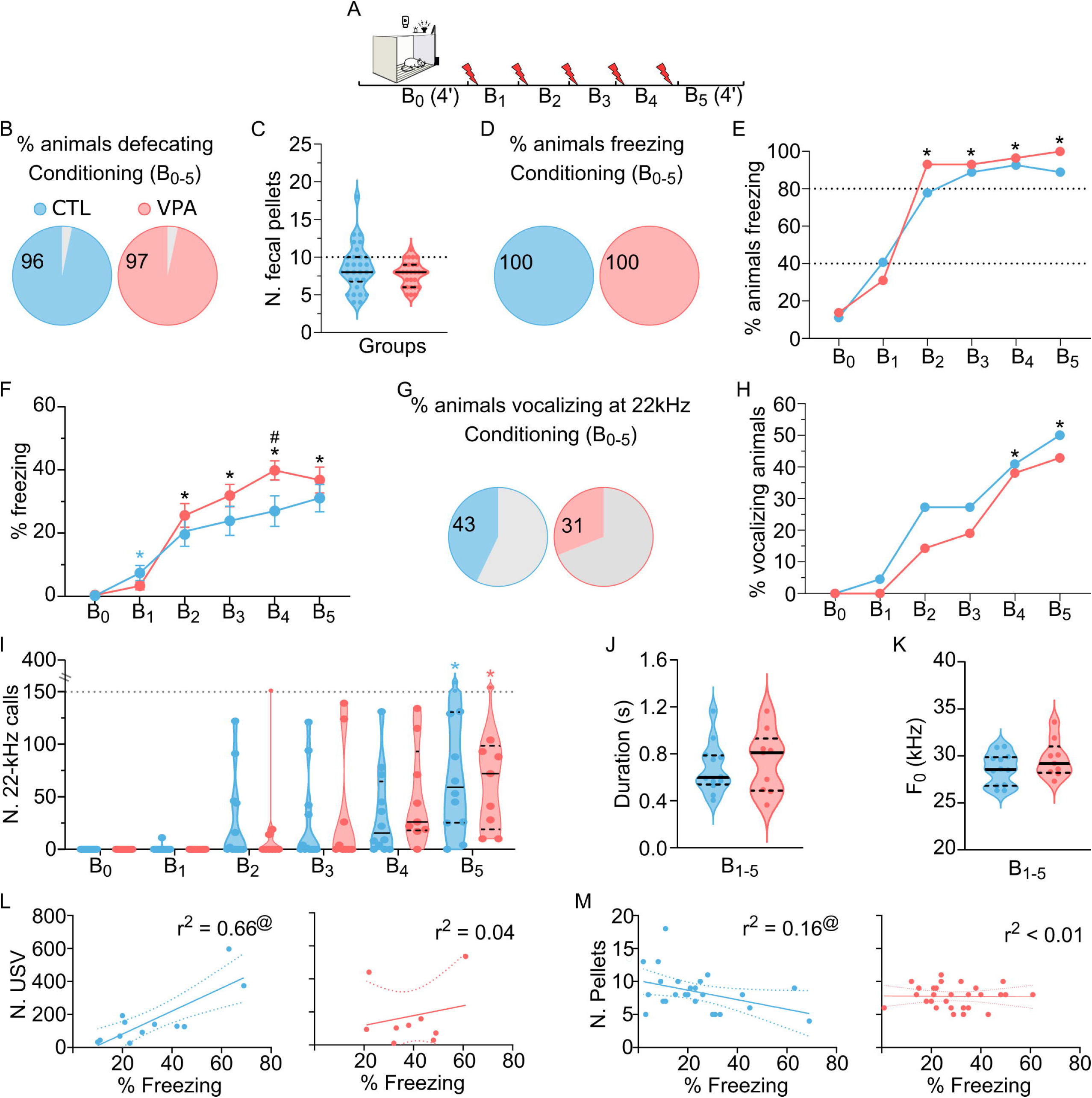
Nociceptive stress response (fear conditioning). (A) Fear conditioning experimental design: Following a 4-min period of free exploration, five electrical shock stimuli were applied. After the last stimulus the animal remained in the chamber for 4-min of free exploration. (B) Percentage of animals releasing fecal pellets. (C) Number of fecal pellets released. (D) Percentage of animals that displayed freezing behavior over all experimental blocks. (E) Percentage of animals that displayed freezing behavior on each experimental block. Dotted lines at 40 and 80% for reference. *p < 0.001. (F) Percentage of time in freezing behavior in each experimental block. B4, ^#^p = 0.046; CTL, *p = 0.010; VPA, *p < 0.001. (G) Percentage of animals that emit USV. ^#^p > 0.050. (H) Percentage of animals that emitted USV on each experimental block. CTL *p ≤ 0.014, VPA *p ≤ 0.026. (I) Number of calls per experimental block emitted by vocalizing animals. CTL *p ≤ 0.003, VPA *p ≤ 0.014. (J) Principal frequency (F0) of calls emitted in blocks B_1-5_. (K) Median call duration in blocks B_1-5_. (L) Correlation analysis between relative freezing behavior and number of long 22-kHz calls emitted by CTL and VPA animals. Each point represents an individual. The trend line indicates the direction and strength of the correlation between behaviors. *p = 0.001. (M) Correlation analysis between relative freezing behavior and number of fecal pellets released by CTL and VPA animals. Each point represents an individual. The trend line indicates the direction and strength of the correlation between behaviors. Panels B, D-E, G-H, Fisher’s exact test. B [N_CTL_ = 31, 15 female, 16 male; N_VPA_ = 30, 14 female, 16 male]; D-E [N_CTL_ = 27; 13 female, 14 male. N_VPA_ = 29; 14 female, 15 male]; G-H [N_CTL_ = 28; 13 female, 15 male. N_VPA_ = 29; 14 female, 15 male]. Panel C Mann-Whitney test [N_CTL_ = 30; 15 female, 15 male. N_VPA_ = 29; 14 female, 15 male]. Panel F, Student-t test and RM one-way ANOVA [N_CTL_ = 27; 13 female, 14 male. N_VPA_ = 29; 14 female, 15 male]. Panel I, Mann-Whitney and Friedman tests [N_CTL_ = 22; 11 female, 11 male. N_VPA_ = 21; 9 female, 12 male]. Panels J-K, Mann-Whitney test [N_CTL_ = 12; 7 female, 5 male. N_VPA_ = 9; 5 female, 4 male]. Panels L, M, Simple linear regression. L [N_CTL_ = 12; 7 female, 5 male. N_VPA_ = 9; 5 female, 4 male]. M [N_CTL_ = 25; 13 female, 12 male. N_VPA_ = 28; 14 female, 14 male]. ^#^, Between-group significant difference. *, Within-group significant difference compared to B0. ^@^, correlation significant difference.

The prevalence of vocalizing animals did not differ significantly between groups, either when analyzed globally or across individual experimental blocks (Fig 3G and 3H).

Both VPA and CTL groups exhibited an increased prevalence of ultrasonic vocalization (USV) emitters from block B4 onward, compared to baseline (B0). Although not statistically significant, a slightly higher proportion of CTL animals vocalized across all blocks relative to the VPA group. Among animals classified as responsive (i.e., those that emitted at least one USV), no between-group differences were found in the total number of 22-kHz calls (Fig 3I). However, a significant within-group increase in call number was observed in both groups at B5. There were no significant differences between groups in the mean duration of the 22-kHz calls or mean fundamental frequency (F_0_; Fig 3J and 3K). Notably, however, vocalizations from VPA-treated animals displayed a bimodal distribution of mean call durations, in contrast to the unimodal pattern observed in CTL. Additionally, a subset of VPA-treated rats (22%; 2/9) produced calls with F_0_ values exceeding the third quartile of the CTL distribution (i.e., >30 kHz).

Given that freezing, 22-kHz ultrasonic vocalizations, and defecation are behavioral indicators of fear, stress, and anxiety, we examined the correlations among these variables. Of the animals tested, 12 out of 27 CTL and 9 out of 29 VPA-treated rats exhibited both freezing behavior and 22-kHz vocalizations, while 26 out of 27 CTL and 28 out of 29 VPA-treated rats displayed freezing behavior and produced fecal pellets. These subsets were included in the respective correlation analyses. In the CTL group, a significant positive correlation was observed between freezing duration and the number of 22-kHz calls, along with a small but significant negative correlation between freezing and the number of fecal pellets released (Fig 3L and 3M). In contrast, no significant correlations between these behavioral measures were found in the VPA group. Finally, analyses of sex differences showed no significant effects on either the prevalence and magnitude of defecation, freezing and vocal response (data not shown).

Memory retrieval was assessed 24 hours after fear conditioning training (Fig S3). Both groups showed a similar proportion of animals exhibiting freezing behavior, with freezing levels significantly elevated relative to baseline (B0). Among responsive animals, freezing levels during the retrieval session did not differ significantly between VPA-treated and CTL rats. To evaluate potential sex differences in freezing behavior, females and males within each treatment group were compared. No significant differences were observed between sexes in either the control or VPA groups (data not shown).

### Prenatal exposure to VPA promotes delayed fear response during emotional contagion

During the emotional contagion test, OBS animals—both VPA-treated and CTL—were exposed to a DEM animal (naive) that either received a series of five-foot shocks or no shocks (Fig 4A). During the experiment, we assessed their behavioral responses by measuring freezing and 22-kHz vocalizations. Our results showed that witnessing DEM animals receiving successive foot shocks engaged more animals of both groups (oC+ and oV+) in freezing behavior. This effect was not observed in OBS witnessing DEM animals that did not received shock (oC- and oV-). Between-group comparisons showed that, at B2 there was a significantly larger engagement in freezing of oC+ animals compared to oC-, as well as of oV+ animals compared to oV-. Notably, in the oV+ group, this elevated freezing response persisted through B3 (Fig 4B). In addition, within-group analysis (compared to B0 baseline), a significantly higher prevalence of freezing behavior was observed in the oC+ group at B2 and B4 and in the oV+ group at B2-B5. We also observed an increase in the relative number of animals expressing freezing behavior at B4-B5 in oC- and oV- groups, but they were not statistically significant.

**Figure 4:**
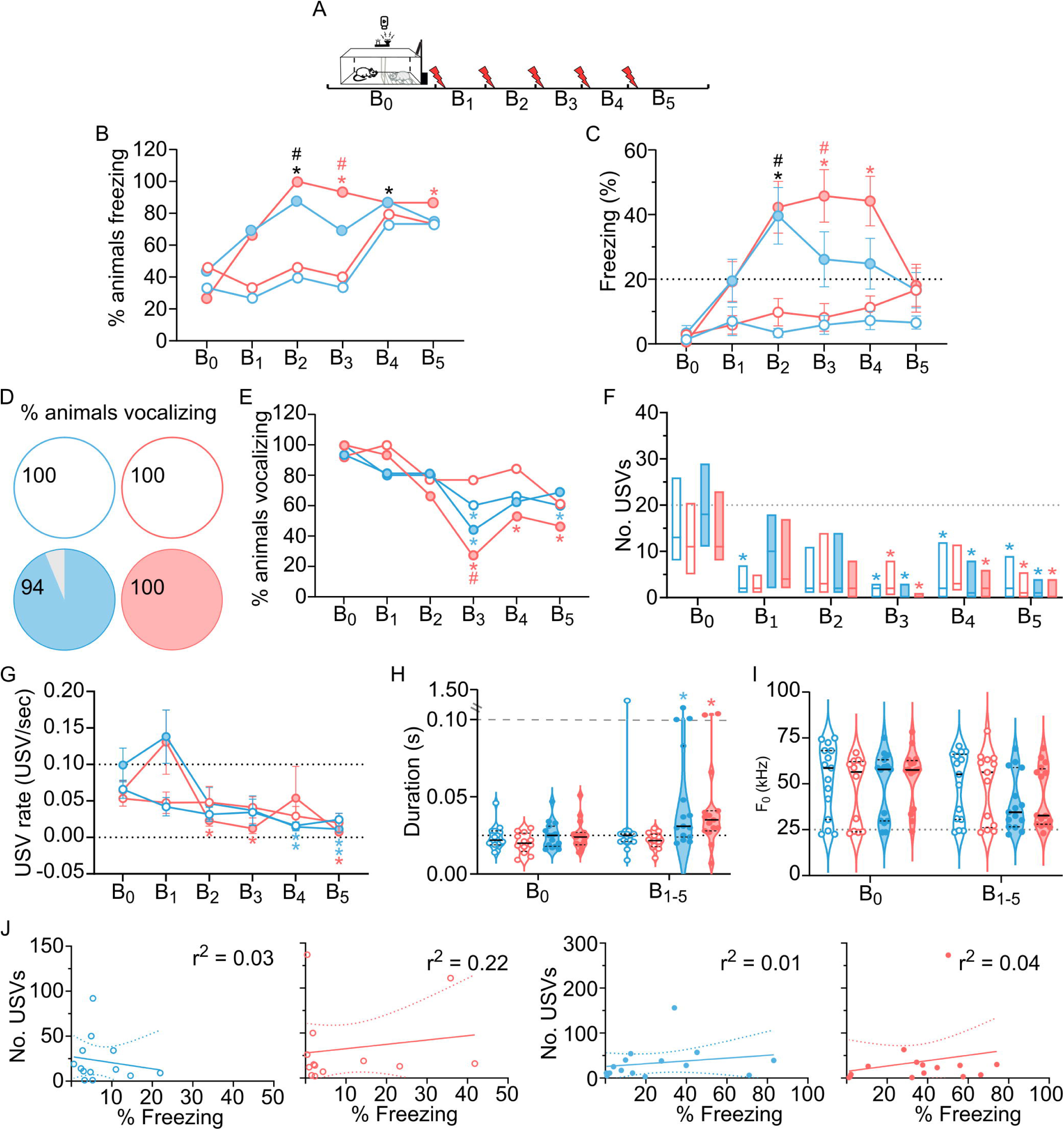
Behavioral analysis during negative social stress (emotional contagion). (A) Experimental design: Following a 4-min period of free exploration, five electrical shock stimuli were applied to the demonstrator. After the last stimulus the animals remained in the chamber for four minutes of free exploration. (B) Percentage of observer animals displaying freezing behavior. (C) Percentage of observer animals displaying freezing behavior on each experimental block. ^#^, p = 0.036; *, p ≤ 0.036. (D) Percentage of time displaying freezing behavior on each experimental block. Dotted line at 20% for reference. ^#^, p ≤ 0.029; *, p ≤ 0.015. (E) Percentage of animals that vocalized long 22-kHz calls over all the experiment. (F) Percentage of animals that vocalized (at all frequencies) on each experimental block. *p = 0.007. (G) Number of calls on each experimental block. Horizontal (dotted and dashed) lines at 20 and 100 calls for reference. Vertical line separate each block. *, p ≤ 0.0306. (H) Call rate on each experimental block. * p < 0.03. (I) Principal frequency of calls over blocks B_1-5_. (J) Call duration over blocks B_1-5_. Dashed line represents 0.1 seconds for reference. (K) Correlation analysis between relative freezing behavior and number of long 22-kHz calls emitted by OBS animals. Each point represents an individual. The trend line indicates the direction and strength of the correlation between behaviors. Panels B,C, E and F, Fisher’s exact test. Panels D and H, two-way ANOVA and RM two-way ANOVA. Panel G, Kruskal-Wallis and Friedman tests. N_oC-_ = 15; 7 female, 8 male. N_oV-_ = 15; 7 female, 8 male. N_oC+_ = 16; 8 female, 8 male. N_oV+_ = 15; 8 female and 7 male) for panels B and C. N_oC-_ = 13; 5 female, 8 male. N_oV-_ = 15; 7 female, 8 male. N_oC+_ = 16; 8 female, 8 male. N_oV+_ = 15; 8 female, 7 male for panel D. N_oC-_ = 15; 7 female, 8 male. N_oV-_ = 13; 6 female, 7 male. N_oC+_ = 15; 8 female, 7 male. N_oV+_ = 15; 8 female, 7 male for panels E-J. N_oC-_ = 13; 5 female, 8 male. N_oV-_ = 13; 6 female, 7 male. N_oC+_ = 15; 8 female, 7 male. N_oV+_ = 15; 8 female, 7 male for panel K.^#^, Between groups: Shock (+) vs. nonShock (-) significant differences. *, Within-group significant difference compared to B0. *, Within-group significant difference compared to Tac. ^&^, female-male within-group difference.

When comparing the freezing behavior of responsive animals (i.e., those exhibiting any freezing) at B2, both oC+ and oV+ rats displayed significantly higher levels of freezing compared to their respective controls (oC- and oV-). Notably, the difference between oV+ and oV- rats persisted through B3 (Fig 4C). Within-group analysis revealed that both oC+ and oV+ animals exhibited increased freezing in response to witnessing repeated shocks administered to their DEM partner, relative to baseline (B0). This increase reached statistical significance starting at B2. Notably, while the heightened freezing in oC+ rats was restricted to B2, oV+ rats maintained an elevated freezing response from B2 through B4.

The prevalence of animals engaging in vocalizations during the experiment (i.e. those animals that emitted at least one 22-kHz or 50-kHz call) was similar between groups, despite a trend to less engagement of animals in oC+ and oV+ groups as more stimuli were applied to DEM (Fig 4D and 4E). Within-group analysis indicated that the oC+, oV+, and oC− groups exhibited a reduction in the proportion of animals vocalizing in response to witnessing their DEM partner receive a shock, relative to B0. This reduction reached statistical significance beginning at B3. Notably, in oC+ rats, the decrease in vocalization was restricted to B3, whereas in oV+ rats, it persisted through B5. Interestingly, the oC- group also showed a significant reduction in vocalization at B5 (Fig. 4E). Changes in the number of vocalizations produced by OBS animals were also observed compared to the baseline, with no differences between groups (Fig 4F). Among responsive animals, a reduction in vocal behavior was observed from block B3 onward in the oC+ and oV+, as well as in oV- groups. In contrast, the oC⁻ animals exhibited a significant decrease as early as block B1, relative to B0. Analysis of vocalization rate (USVs per minute) revealed that oV+ animals exhibited a consistent reduction during sessions B2, B3, and B5. In contrast, both oC+ and oC- groups showed a decline in vocal behavior only from session B4 onward. The oV- group maintained a stable vocalization rate across all sessions. Notably, both oC+ and oV+ animals displayed a peak in vocalization rate during session B1, followed by a progressive decline in subsequent blocks. (Fig 4G).

When basic bioacoustic properties (duration and f_0_) of the ultrasonic calls were analyzed, we noticed that calls produced by oC+ and oV+ animals had longer durations at blocks B1-B5 when compared to calls produced at B0 (Fig 4H). However, no differences were observed in f_0_ either across blocks or between groups (Fig 4I). Correlation analysis between freezing and vocal production released during the emotional contagion showed no significant differences (Fig 4J). No sex-related differences were observed in the susceptibility and magnitude of the freezing and vocal responses (data not shown).

Considering that the behavioral responses displayed by OBS animals should depend on the behavior of DEM rats, we also quantified freezing and vocalizations produced by these animals. Exposure to shock significantly increased prevalence of displaying freezing compared to non-shocked controls (dV− and dC−), regardless of pairing with VPA-treated or CTL counterparts. (Fig S4A-B). No significant differences were observed between the dV+ and dC+ groups or between the dV− and dC− groups. Among responsive animals, both dC+ and dV+ groups exhibited a significant increase in freezing behavior following the delivery of shock stimuli, as expected (Fig S4C). Compared to B0, dC+ and dV+ increased freezing from B1 to a plateau between B2-B5. dV- and dC- animals showed no changes in freezing behavior relative to B0 levels. Importantly, no differences were observed in freezing between either dC+ vs. dV+ or dC- vs. dV- (p > 0.05).

Among dV+ and dC+ animals, there was a high prevalence of vocalizing animals compared to dV- and dC- rats (Fig S4D). The vocal engagement across experimental blocks revealed (1) no group differences between dV+ vs. dC+ or between dV- vs. dC-; (2) significant differences between dV+ vs. dV- (with onset at B2) and between dC+ vs. dC- (with onset at B1) and (3) an increase relative to B0 levels in dV+ and in dC+ (from B2 onwards; Fig S4E). dV- and dC- animals showed no changes in vocal engagement relative to B0 levels.

DEM rats exhibited robust engagement in fear-related ultrasonic vocalizations (USVs) following foot shock exposure during the emotional contagion experiment. No significant differences in the number of USVs were observed between dV+ and dC+ groups; however, both groups demonstrated a marked increase in USV emissions from day B4 onward relative to their respective controls (dC- and dV-; Fig S4F).Besides, significant vocal production increased was observed in dV+ (at B2-B5), dC+ (at B2, B4-B5), as well as dV- (at B1) animals, compared to B0. No significant within-group changes were observed in calls emitted by dC- rats. Bioacoustic properties such as call duration and f_0_ did not change between dV+ and dC+ (Fig S4G and S4H). Owing to the low number of vocalizations observed in dV- and dC- animals (median of 3 per block), their acoustic properties were not analyzed.

## Discussion

Animal models remain essential for elucidating the mechanisms underlying emotional dysregulation in neurodevelopmental disorders and for developing therapeutic strategies. In the present study, we systematically examined the stress responsiveness across multiple behavioral paradigms and the emotional contagion response in a rat model of autism induced by the prenatal exposure to VPA.

Unfamiliar handling in rodents is known to elicit stress-related behaviors such as avoidance and increased defecation, which typically attenuate with repeated exposure to the experimenter (39–41). In the present study, the greater number of VPA-treated animals releasing fecal pellets, as well as the greater number of fecal pellets released during the first experimental days, compared do CTL, suggest an increased susceptibility to stress responses to innocuous stimuli. Furthermore, although both VPA-treated and CTL animals showed a reduction in fecal pellet production after four days of handling, suggesting successful habituation responses in both groups, the delayed reduction observed in VPA rats indicates a slower adaptation to nonthreatening stimuli, consistent with reports of atypical sensory processing, stress management, and impaired desensitization in individuals with ASD. Evidence from human studies consistently demonstrate altered responses to tactile stimuli in ASD, characterized by both hyper- and hypo-behavioral reactivity, but with hyper-reactivity being the most consistently observed phenotype (42–49). These behavioral manifestations suggest autonomic nervous system (ANS) dysregulation. This alteration is accompanied by elevated baseline electrodermal activity and heart rate, corroborating the dysregulation of the ANS (13–15). A recent study examining ANS responses to affective touch (e.g., gentle stroking of hairy skin) in children reported smaller autonomic reactivity in those with ASD (13). Similarly, the increased defecation response of VPA rats during early handling sessions may reflect analogous autonomic dysregulation and an elevated susceptibility for stress responses to innocuous stimuli.

Although a greater proportion of VPA-treated animals exhibited defecation in response to electro-tactile stimulation compared to controls, suggesting increased susceptibility, neither the freezing behavior nor the number of fecal pellets differed significantly between groups, indicating comparable magnitudes of stress responses. These findings are consistent with our observations of tactile and nociceptive thresholds and align with previous studies demonstrating preserved nociceptive thresholds to electrical stimulation in VPA-exposed rats (50). Notably, prior investigations into tactile sensitivity in VPA-treated animals have consistently reported increased responsiveness to mechanical allodynia and diminished sensitivity to thermal nociceptive stimuli (35,36,51–55).

During the fear conditioning paradigm, VPA and CTL animals were similarly susceptible to stress-induced defecation, freezing, and 22-kHz vocalization behaviors, following a similar time-course in both groups. Although studies examining the dynamics of the freezing response in VPA-treated animals remain limited, our findings are consistent with a previous report showing increased shock-induced freezing in VPA rats relative to CTL controls (56). In contrast, other studies assessing overall freezing across the entire experimental session have reported preserved freezing behavior in VPA-treated animals (36,50,57). These discrepancies may arise from variations in experimental design, strain-specific factors, or developmental stage differences. Further analysis of our data indicated a delayed onset of freezing behavior in the VPA group, potentially reflecting impairments in the integration or coordination of stress-related emotional responses, including sensory processing, decision-making, motor planning and execution, or disruptions in emotional and motivational regulation. This delayed response, alongside preserved vocalization, reinforces the hypothesis of altered integration or coordination of emotional processing under stress. Moreover, the observed negative correlation between freezing behavior and fecal pellet release—along with the progressive increase in freezing observed in CTL animals—suggests that defecation predominates at lower stress intensities, whereas freezing becomes more prominent as stress levels escalate. The lack of this correlation in VPA-treated rats provides additional evidence for a disruption in the integration of physiological responses associated with emotional stress.

During fear retrieval, behavioral responses were limited, particularly for vocalizations and defecation, restricting the analysis to freezing behavior. No group differences were observed in freezing magnitude or incidence. These findings are consistent with prior reports indicating preserved retrieval responses in VPA rats and further suggest that acute associative memory of the conditioned stimulus is intact in this model.

Nociceptive responses in individuals with autism spectrum disorder (ASD) vary across developmental stages. Human studies have shown that children with ASD often exhibit hypersensitivity to pressure and cold pain, while adolescents tend to display thermal pain thresholds within the normative range. In adults, thermal pain thresholds are preserved, but pain ratings are elevated and more variable (58–65). These developmental discrepancies in pain sensitivity further underscore the complexity and heterogeneity of sensory and stress-related phenotypes in ASD. Our findings—demonstrating preserved nociceptive sensitivity but altered stress habituation in adolescent rats—parallel this developmental trajectory and suggest translational relevance for the VPA model in this domain.

In the emotional contagion paradigm, both CTL and VPA observer rats exhibited freezing and 22-kHz vocalizations in response to a distressed conspecific, consistent with prior findings in CTL animals (66,67). However, VPA observers displayed prolonged freezing and delayed habituation, suggestive of increased sensitivity to social distress or impairments in social buffering mechanisms. Notably, demonstrator animals showed comparable freezing behavior across groups (Fig. S3B–C) and only minor differences in vocalizations (Fig. S3D–F), indicating that the sensory cues available to observers were largely equivalent. Additionally, the acoustic properties of demonstrator calls were similar across conditions, supporting the conclusion that CTL and VPA observers were exposed to comparable emotional signals (Fig. S3G–H). Together, these results indicate that VPA-treated animals exhibit altered habituation and heightened responsiveness to social distress, reflecting a dysregulated emotional contagion response. This phenotype underscores the utility of the VPA model for investigating stress reactivity and social emotion regulation deficits relevant to ASD.

While emotional contagion has been well-characterized in various rodent strains, to our knowledge, no previous studies have examined this phenomenon in a rodent model of autism. Our findings thus provide novel evidence that, despite their established social deficits, juvenile VPA animals still perceive social stressors as salient, though more challenging to process. In addition, the preserved sensitivity to social cues but fail to downregulate stress responses following social exposure of VPA-treated animals results are consistent with human data, where both children with ASD and TD peers show increased cortisol levels in response to social interaction. However, while TD children demonstrate cortisol habituation by the end of the exposure, this attenuation is absent in children with ASD. Additionally, among children exhibiting the highest cortisol reactivity— including both ASD and TD groups—those with ASD engaged in lower levels of social interaction. Complementary findings from the same research group further report that elevated cortisol responses to social challenges are positively associated with increased daily stress and sensory processing difficulties (68–71). This pattern underscores the translational value of the VPA model for investigating stress regulation abnormalities in ASD. By capturing both behavioral and physiological features of impaired social stress adaptation, the model offers an important platform for elucidating the underlying neurobiological mechanisms and for identifying potential targets for therapeutic intervention.

Notably, across all experimental paradigms, VPA-treated animals consistently exhibited a trend toward increased involuntary responses (defecation and freezing) and slightly reduced voluntary vocalizations relative to control animals, even in the absence of statistically significant group differences.

Sex differences in stress responsiveness were not observed across any of the para- digms employed. This aligns with previous research demonstrating that juvenile rats ex- posed to VPA do not exhibit significant sex-based differences in social play behavior (72). These findings are consistent with a human study in which VPA-exposed ASD children have a much reduced autism male bias (1.2-2.4:1) compared to no VPA-exposed ASD children (3.3:1) (73,74). One possible explanation for the attenuated sex differences is that VPA acts as a neurodevelopmental risk factor whose effects may override or bypass sex- dependent protective mechanisms. Alternatively, it has been proposed that females with ASD may engage in compensatory social behaviors that mask symptomatology, contributing to diagnostic underrepresentation and an artificially reduced prevalence (75–78). The absence of sex effects in our study may also reflect limitations in the sensitivity of the employed behavioral assays to detect subtle sex-dependent phenotypic variations or may result from the homogeneity of the VPA-induced phenotype across sexes. To better elucidate sex-specific mechanisms, future studies should incorporate hormonal profiling or cir- cuit-level analyses and more nuanced behavioral assessments designed to detect subtle, sex-specific traits.

Finally, we would like to identify some limitations of the present study that include the lack of assessment of hearing as they would be particularly important to control for individuality in hearing-dependent fear responses, as well as the lack of measurements of corticosterone levels, which would have been valuable to control for eventual stress-related HPA dysfunctions VPA-treated animals. Considering these limitations will provide future experiments with important directions for improvement.

## Conclusion

VPA-treated rats exhibited altered stress and fear-related behaviors across multiple paradigms. Compared to controls, they showed heightened and prolonged physiological responses to handling and electro-tactile stimulation, as well as delayed and exaggerated freezing during fear conditioning. The typical correlations among fear-related behaviors observed in control animals were disrupted in the VPA group, indicating a decoupling of behavioral stress markers. During emotional contagion, VPA-treated observers demonstrated more sustained freezing and earlier reductions in vocalization when witnessing a conspecific in distress, suggesting altered emotional processing. These findings support a profile of increased emotional reactivity and impaired integration of fear-related responses in VPA-treated animals. Collectively, these findings suggest that the VPA model provides a valid platform for investigating the neurobiological mechanisms of stress susceptibility and social dysfunction in ASD.

## Acknowledgments

We are grateful to Renan Araújo de Lima, Ana Luiza Campos Vila Real and Juliana Alves Brandão Medeiros de Sousa for their technical assistance with experimental apparatus design, behavioral experiments and histological procedures.

## Author’s contribution

Author DH: Conceptualization, data curation, formal analysis, investigation, methodology, project administration, supervision; validation, visualization, writing – original draft preparation, and writing – review and editing.

Author ALD: Formal analysis, and writing – review and editing.

Author RNRP: Conceptualization, formal analysis, funding acquisition, methodology, resources, writing – review and editing.

## Supporting informations

**S1_Fig: USV power.** (A) Maternal separation (MS) at P08 schematic illustration and graphic with USVs mean power by animal. ^#^p < 0.0001. (B) Maternal separation (MS) at P14 schematic illustration and graphic with USVs mean power by animal. ^#^p < 0.0001. (C) Fear conditioning (FC) at P35-38 schematic illustration and graphic with USVs mean power by animal. Dotted line represents the lower power of USV emitted in the microphone chamber. Dashed line represents the higher power of USV emitted in the opposite chamber. Panels A, B and C, Mann-Whitney test. N opposite = 41. N microphone = 35 for panel A. N opposite = 55. N microphone = 69 for panel B. N opposite = 153. N microphone = 153 for panel C. ^#^, between-group significant differences.

**S2_Fig: Tactile and nociceptive thresholds.** (A) Tactile and nociceptive thresholds. ^#^p > 0.050; *p < 0.001. (B) Female and male tactile thresholds. ^&^p = 0.007. Panel A, paired Student-t test [N_CTL_ = 30; 15 female, 15 male. N_VPA_ = 28; 15 female, 13 male]. Panel B, Mann-Whitney [N_CTL_ = 30; 15 female, 15 male. N_VPA_ = 28; 15 female, 13 male]. ^&^, between-group female/male significant differences.

**S3_Fig: Memory retrieval.** (A) Experimental design: the rats were placed in the middle of the apparatus facing one of the metal walls. After five minutes, animals return for its home cage. (B) Percentage of animals that displayed freezing behavior over the experiment. (C) Percentage of animals that present freezing behavior throughout the experiment. *p = 0.0003. Panel B, Fisher’s exact test. Panel C, Student-t and paired Student-t tests. N_CTL_ = 27; 13 female, 14 male. N_VPA_ = 28; 15 female, 13 male. *, within-group significant differences.

**S4_Fig: Emotional contagion DEM.** (A) Emotional contagion experimental design: Following a 4-min period of free exploration, five electrical shock stimuli were applied to the demonstrator. After the last stimulus the animals remained in the chamber for four minutes of free exploration. (B) Percentage of demonstrator animals that present freezing behavior on each experimental block. CTL ^#^p ≤ 0.011; VPA ^#^p ≤ 0.021; dC+ *p < 0.001; dV+ *p ≤ 0.012. (C) Demonstrator’s freezing behavior percentage in each experimental block. CTL ^#^ p ≤ 0.008; VPA ^#^p < 0.001; dC+ *p < 0.001; dV+ *p ≤ 0.022. (D) Percentage of animals that emitted USVs throughout the experiment (B0-5). CTL ^#^p < 0.001; VPA ^#^ p ≤ 0.011. (E) Percentage of animals that emitted USVs on each experimental block. CTL ^#^p ≤ 0.002; VPA ^#^p ≤ 0.006; dC+ *p ≤ 0.042; dV+ *p ≤ 0.027. (F) Number of USVs on each experimental block. CTL ^#^ p ≤ 0.017; VPA ^#^ p ≤ 0.024; dV- *p = 0.017; dC+ *p ≤ 0.001; dV+ *p ≤ 0.042. (G) USVs duration emitted in blocks B_1-5_. (H) USVs principal frequency emitted in blocks B_1-5_. ^#^, Between groups: Shock (+) vs. nonShock (-) significant differences. *, Within-group significant difference compared to B0.

