## Supplementary figures and images for "Altered Stress and Fear Responses in the VPA Rat Model of Autism: Behavioral Dissociation Across Tactile, Nociceptive, and Social Contexts"

### Figure S1

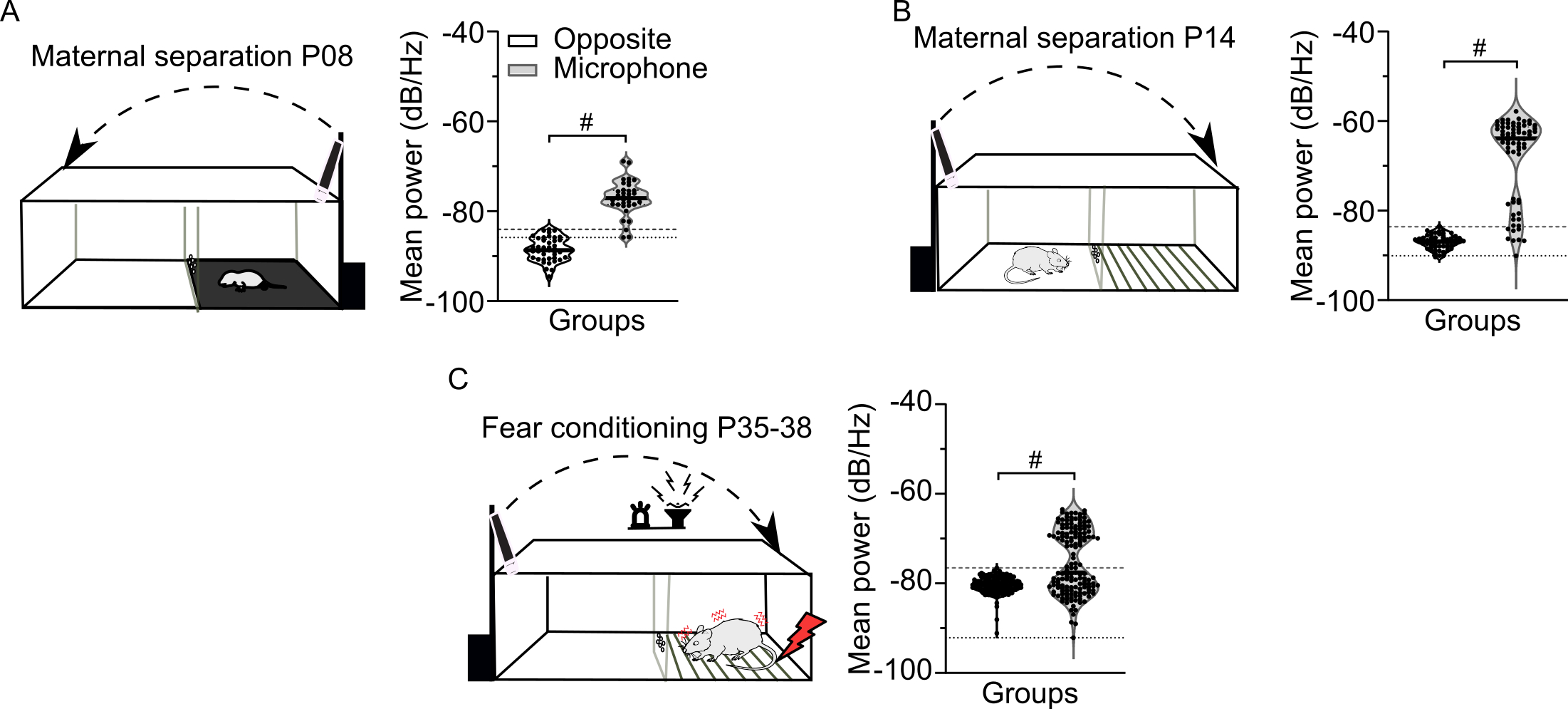

### Figure S2

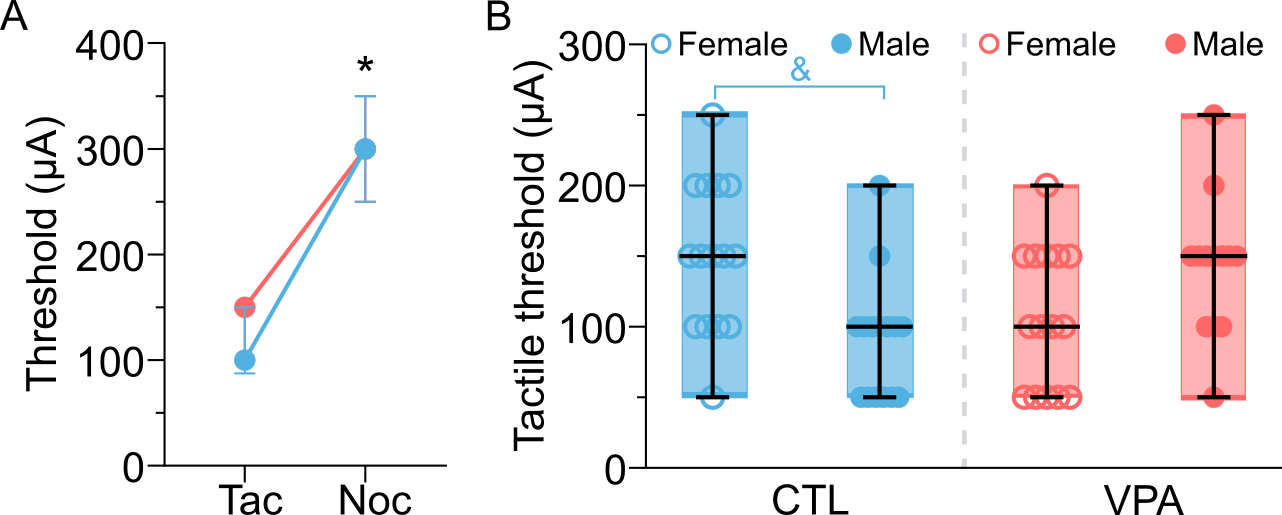

### Figure S3

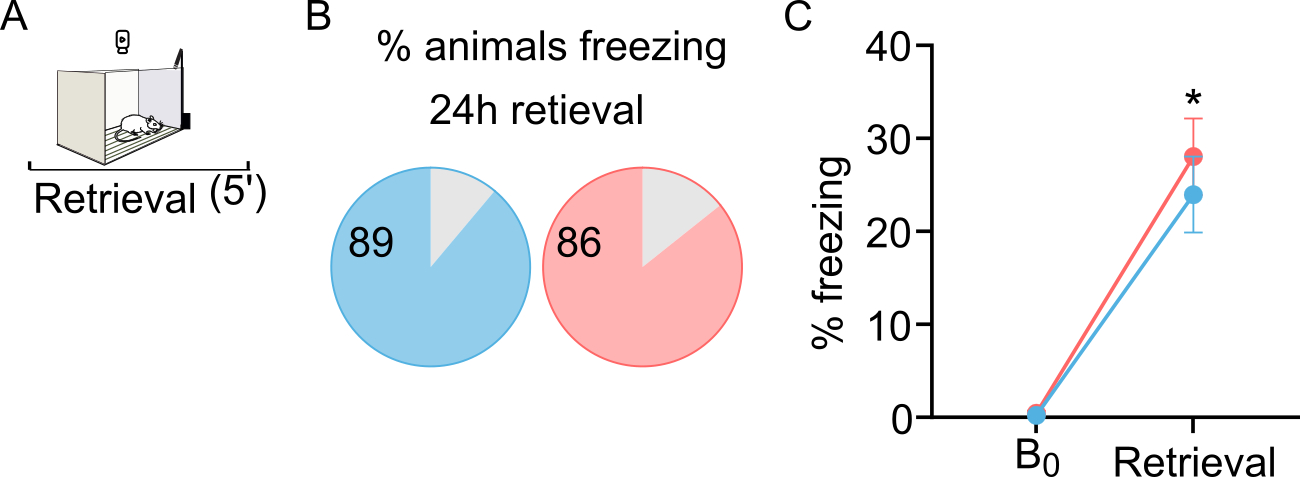

### Figure S4

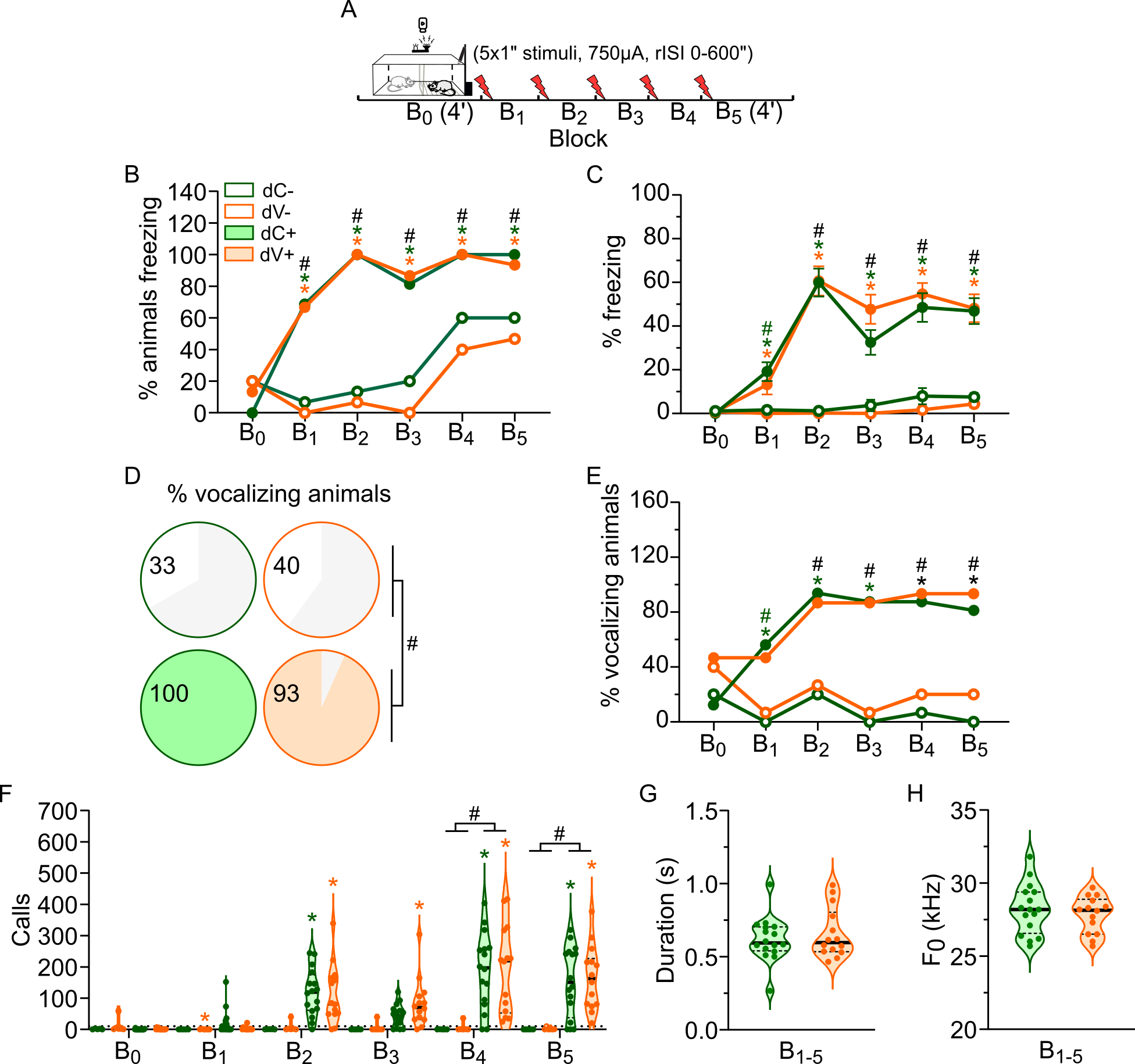
